# One-Shot Random Forest Model Calibration for Hand Gesture Decoding

**DOI:** 10.1101/2023.07.21.550033

**Authors:** Xinyu Jiang, Chenfei Ma, Kianoush Nazarpour

**Affiliations:** School of Informatics, The University of Edinburgh, Edinburgh, United Kingdom

**Keywords:** myoelectric control, electromyography, random forest

## Abstract

**Objective:** Most existing machine learning models for myoelectric control require a large amount of data to learn user-specific characteristics of the electromyographic (EMG) signals, which is burdensome. Our objective is to develop an approach to enable the calibration of a pre-trained model with minimal data from a new myoelectric user.

**Approach:** We trained a random forest model with EMG data from 20 people collected during the performance of multiple hand grips. To adapt the decision rules for a new user, first, the branches of the pre-trained decision trees were pruned using the validation data from the new user. Then new decision trees trained merely with data from the new user were appended to the pruned pre-trained model.

**Results:** Real-time myoelectric experiments with 18 participants over two days demonstrated the improved accuracy of the proposed approach when compared to benchmark user-specific random forest and the linear discriminant analysis models. Furthermore, the random forest model that was calibrated on day one for a new participant yielded significantly higher accuracy on day two, when compared to the benchmark approaches, which reflects the robustness of the proposed approach.

**Significance:** The proposed model calibration procedure is completely source-free, that is, once the base model is pre-trained, no access to the source data from the original 20 people is required. Our work promotes the use of efficient, explainable, and simple models for myoelectric control.

## 1. Introduction

Many myoelectric control systems use machine learning to map the electromyographic (EMG) signals [1, 2, 3, 4, 5] to control commands for human-machine interfaces, e.g. prosthesis [6, 7, 8, 9, 10, 11, 12] and virtual keyboards [13, 14]. Most modern myoelectric control machine learning models require a large amount of data from a user to learn a bespoke and user-specific map [7, 15, 16]. In real-life settings, machine learning models that are flexible and require minimal data from a new user are preferable [17]. Additionally, such models are expected to be robust in long-term (multi-day) applications without the need for frequent re-calibration.

Domain adaptation minimises the distribution mismatch between training data (source domain) and testing data (target domain) [18]. In myoelectric control applications, this technique can be used for a model to learn extra knowledge from source data collected from other subjects, so that the demand for data collection from a new target user is reduced. For instance, Vidovic et al. [19] employed a covariate shift adaptation algorithm to align the statistical metrics (e.g., the mean value and covariance) of source and target domains. These statistical metrics informed the training of a linear discriminant analysis (LDA) model. Wang et al. [20] developed a multi-user myoelectric control model using discriminative canonical correlation analysis (CCA) by jointly projecting the feature sets for source and target domains into a low-dimensional, uniform-style subspace. Xue et al. [21] integrated CCA and the optimal transport (OT) mechanism, termed CCA-OT, to further reduce the distribution discrepancies between the source and target domains. Jiang et al. [22] proposed a correlation-based data weighting (CORW) scheme to measure the level of feature distribution shift between the target testing user and each training user and then assign different weights to data from different training subjects accordingly. One common disadvantage of the above approaches is the need to access the data in the source domain. EMG data contain diverse private and sometimes sensitive information, e.g., the status of neurological diseases [23] or personal identity [24]. Therefore, in most practical cases, the source EMG data should be strictly protected, due to regulations such as the European General Data Protection Regulation (GDPR) and user concerns.

Large pre-trained models have proven effective in many machine learning tasks and shown their impressively strong generalisation capability, e.g., the Generative Pre-trained Transformer 4 (GPT-4) [25] and the segment anything model (SAM) [26] in computer vision (CV). However, such large pre-trained models are mainly built on differentiable and parameterised deep neural networks. The pre-training of such large models with back-propagation relies heavily on extremely large datasets and significant computing infrastructure. Despite their excellent performance, such pre-trained models have a highly complex architecture and are used as a black-box module. In myoelectric control applications, large pre-trained models are not available up to date in the academic and clinical sectors, mainly because collecting an extremely large and labelled dataset is expensive and time-consuming. Additionally, for most applications of myoelectric control, the model should be implemented in a portable and wearable system with limited computing and battery resources. Moreover, the employed model is expected to be explainable [27, 28, 29]. Therefore, we asked: “Is it possible to create a pre-trained myoelectric control model that can be tuned using a small dataset for a new user, is explainable, and can classify the EMG signals in real-time with limited computing resources, e.g., memory?”

Standard, off-the-shelf random forest (RF) models have not been generally viewed as the most powerful model for myoelectric control [30]. However, RF models have proved effective in dealing with problems with small sample sizes [31]. They can capture complex nonlinear relations between samples. They exhibit robustness on outliers and they can be easily parallelised for computational efficiency [32, 33]. Additionally, the decision tree model, which is the base learner in RF, is recognised as one of the most explainable models [34, 29]. The above merits motivated us to explore new possibilities with RF models and its potentials for myoelectric control applications.

We propose a myoelectric control model with a user-generic RF model that is pre-trained using data from 20 users. When a new (i.e., 21st) user is introduced, we use new EMG data with only 1s signal duration for each gesture to fine-tune the pre-trained RF model, upon a series of tree pruning [35], branch grafting [36, 37], and tree appending procedures. We show that our approach improves model accuracy and across-day generalisability significantly. Crucially, the proposed model tuning is completely source-free, that is we do not need the source data (from the original 20 participants) to tune the pre-trained model for a new user. Both offline validations and the real-time implementation demonstrated the superiority of our method compared with the standard user-specific RF and the benchmark LDA models. Overall, the contributions of our work are summarised below:

1. We provide a transfer learning mechanism in EMG pattern recognition using simple, easily parallelisable, computationally efficient, and explainable RF, which is a practical alternative solution in mobile applications with limited computation resources.
2. The proposed RF calibration method also achieves better across-day generalisability compared with efficient standard LDA and RF models.
3. The proposed RF calibration method is source-free, satisfying the data privacy regulations in practice.
4. This is also the first attempt in myoelectric control area to develop one-shot calibration methods on simple and efficient RF models. The promising results can inspire more follow-up studies in the new research track.

## 2. Methods

### 2.1. Ethical approval

All subjects signed an informed consent form which was approved by the local ethics committee at the University of Edinburgh (reference number: 2019/89177). We conducted two experiments.

### 2.2. Experiment 1

Experiment 1 was offline. We collected EMG data from 20 able-bodied subjects (12 males, 8 females, age range: 22–43). Eight electrodes were equally spaced across the circumference of the forearm, as presented in Figure 1. The EMG data were collected using Delsys Trigno sensors (Delsys, Inc.), with a sampling rate of 2000 Hz and bandpass filter of 10–500 Hz. Each subject performed six hand gestures, as presented in Figure 1. For each gesture, 10 repetitions were performed in 10 successive trials, each six seconds long. In each trial, subjects were required to mimic an instructed target hand gesture and then hold the gesture until the trial ended. Signals recorded in the first two seconds of each trial were removed to get rid of the transient period. The EMG signals within the last four seconds were retained. There was a five-second resting period between trials. Data from Experiment 1 was used to pre-train the RF model.

**Figure 1.**
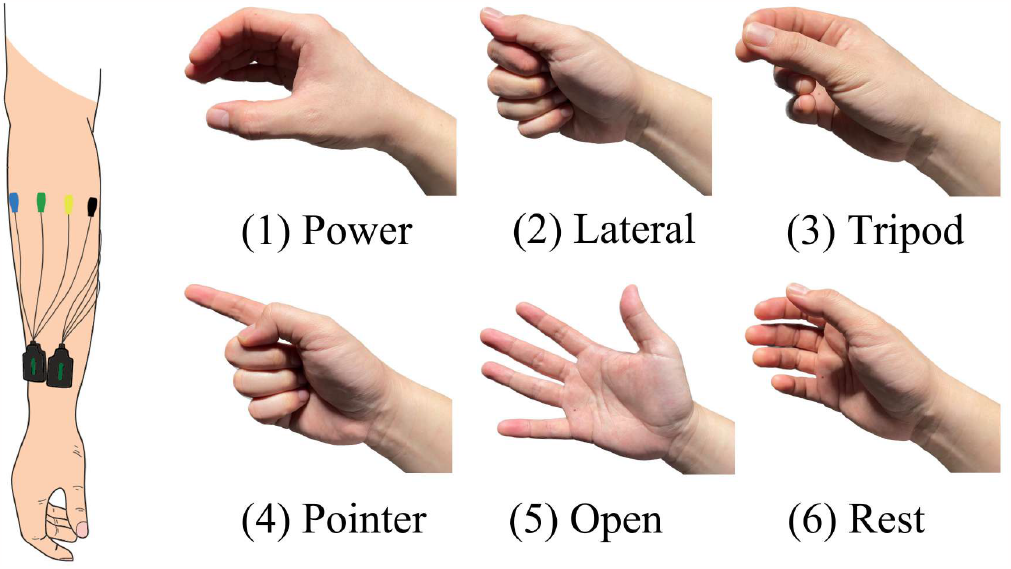
Electrode positions and involved gestures in our experiment.

### 2.3. Experiment 2

Experiment 2 was a real-time myoelectric control experiment. We recruited 18 *new* able-bodied subjects (11 males, 7 females, age range: 22–28). Experiment 2 was conducted on two successive days. On the first day, the experiment consisted of two sections, namely, calibration and testing. In the calibration section, subjects were asked to perform each hand gesture only once in a 2s trial. Each two-second trial afforded the participants one second to shape their hand so that they could hold the gesture reliably during the second one-second period. Only the data during the latter period was used for model tuning, or training a user-specific model from scratch (as a benchmark).

The testing section comprised five blocks. In each block, five repetitions of each gesture were instructed in a pseudo-randomised way. Subject had two seconds and five minutes for rest between trials and between blocks, respectively. On the second day, subjects started the testing section without any further tuning. Electrodes were replaced on day 2 according to the electrode positions marked on day 1.

### 2.4. Feature Extraction

In both experiments, we used a window of 200 ms, sliding at 100 ms, for feature extraction. Ten features were extracted, which described the EMG data from three complementary aspects. First, the mean absolute value (MAV), waveform length (WL), zero crossings (ZC) and slope sign changes (SSC) of the EMG signals, combined with root mean square (RMS), were used as energy descriptors [38, 39, 40]. Second, the skewness [41, 42, 43] of the EMG signals was extracted as a distribution descriptor. Third, the mean frequency (MNF), median frequency (MDF), peak frequency (PKF), and variance of central frequency (VCF) were extracted as spectrum descriptors [44]. We will demonstrate the complementary roles of these features through an ablation experiment in the Results Section.

#### Algorithm 1

Decision Tree Pruning

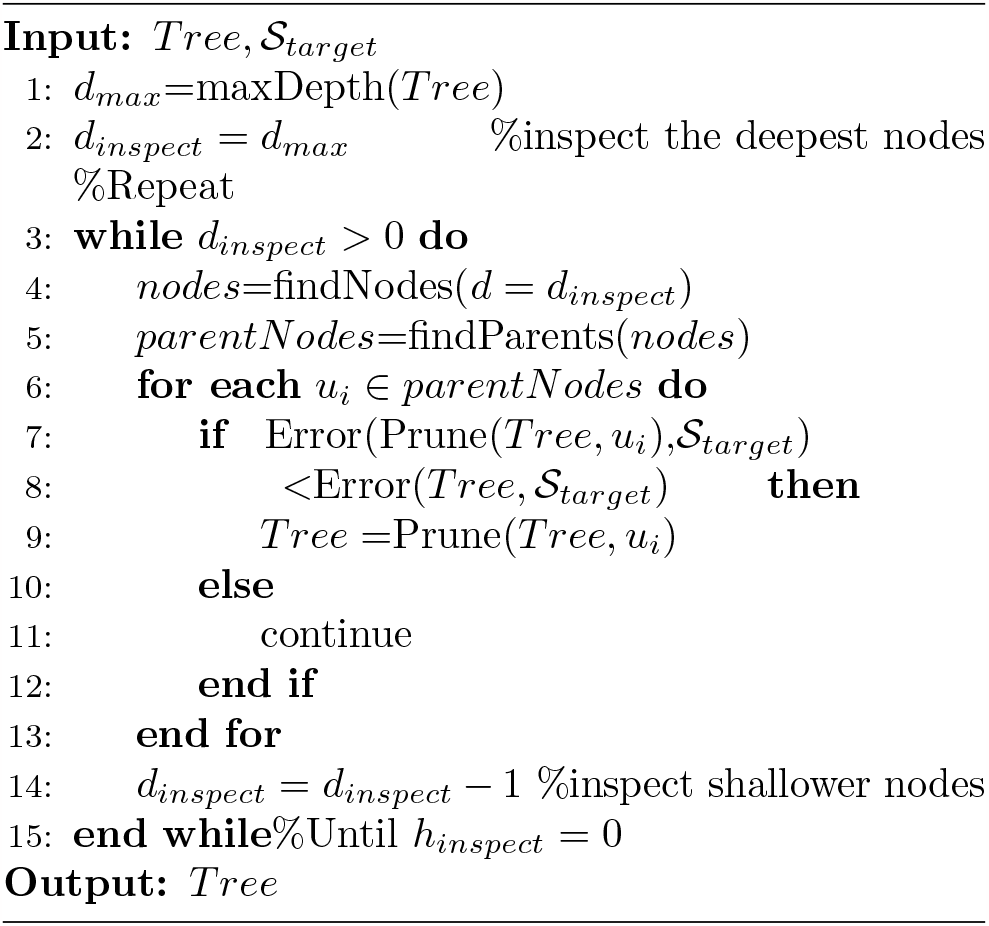

#### Algorithm 2

Decision Tree Grafting

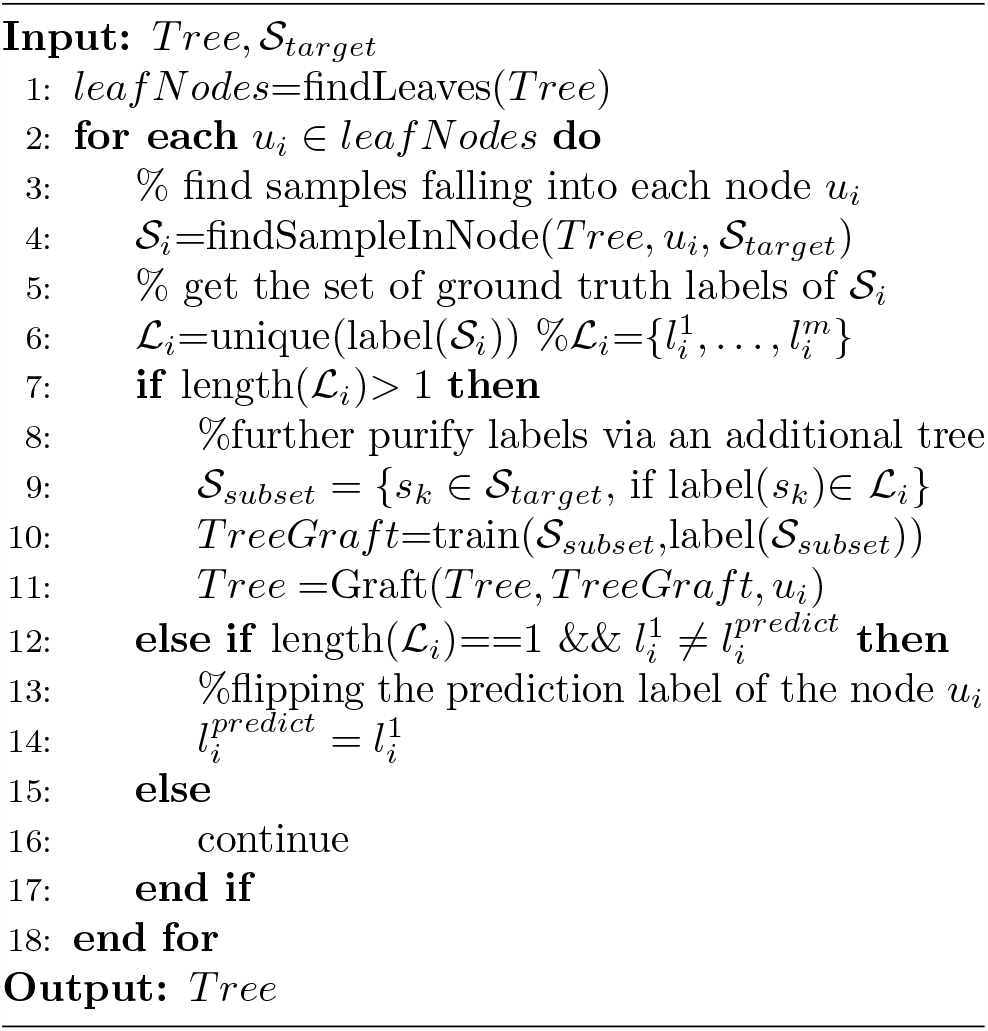

### 2.5. The Proposed RF Model

#### 2.5.1. Pre-training the RF model

Data from Experiment 1 was used for pre-training an RF model. The model had 200 decision trees. A subject-wise feature normalisation was performed on the pre-training dataset. Specifically, the extracted features from each subject were normalised separately to a mean value of zero and a standard deviation of one to align the basic statistical metrics of feature distributions from different subjects. The allocation of pre-training and testing subjects varies with different validation settings and will be described later.

#### 2.5.2. Pruning and Grafting Decision Trees

Given a pre-trained RF model, the follow-up calibration procedure involved three strategies, namely, pruning, grafting, and appending decision trees, as presented in Figure 2. Pruning is a widely used strategy to simplify a model and at the same time improve its generalisability [45]. Grafting, that is, growing new branches on a leaf node to further specify the decision rule. Compared to pruning, grafting has attracted less attention in previous studies, but both the pruning [35], grafting [46, 37] or the combination of both [36] have been proven effective in fine-tuning a decision tree in different application scenarios. In this work, we applied both strategies and compared their performances in myoelectric control applications to provide a comprehensive benchmark for future research. Either pruning or grafting can contribute to a satisfactory performance.

**Figure 2.**
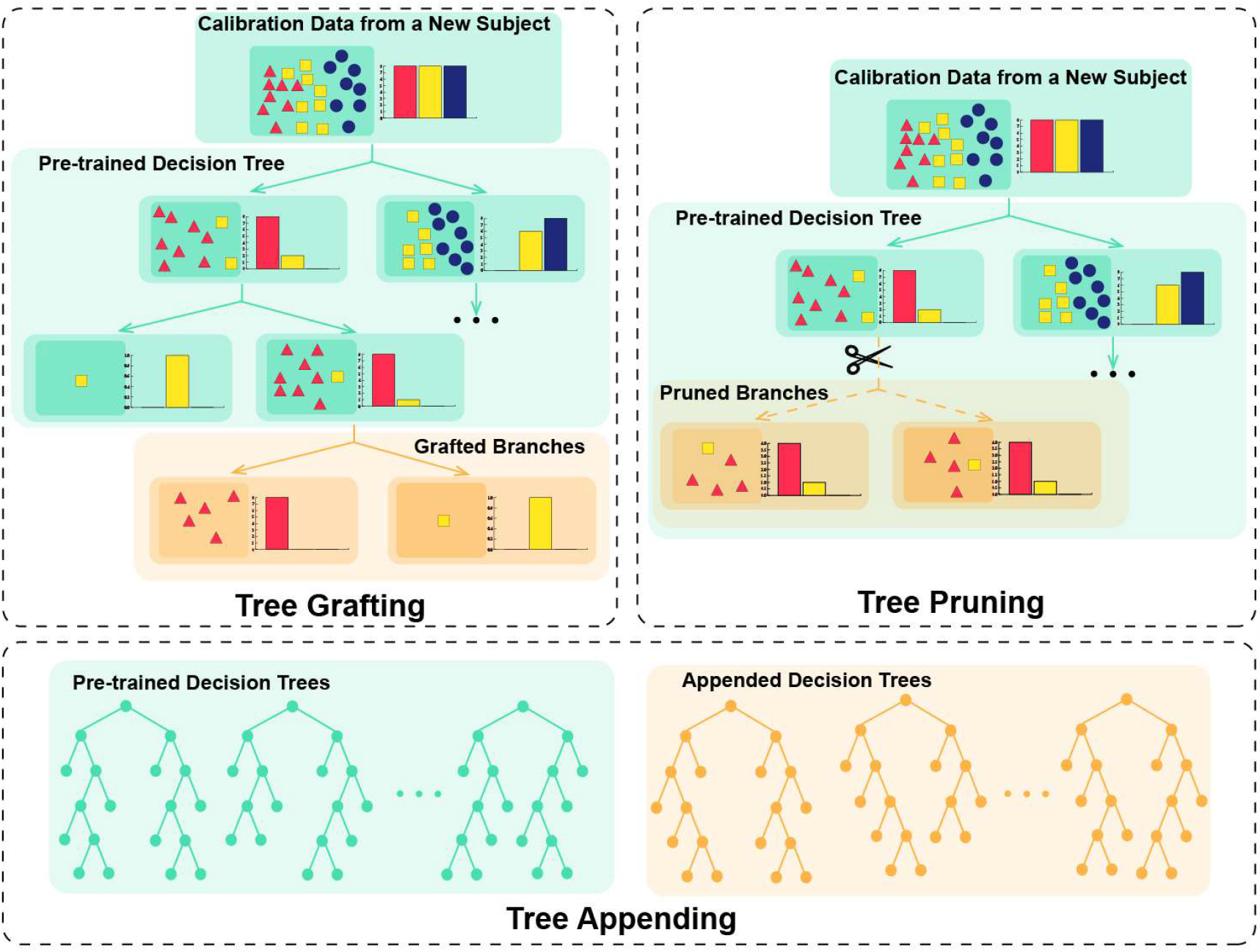
Tree grafting, tree pruning, and tree appending for RF model tuning.

For pruning, we applied a bottom-up strategy. Let’s assume a pre-trained decision tree *Tree* with a node *u*_*i*_ at depth *d*_*i*_, where the depth of the root node is zero and the depth of a leaf node with the longest decision path is *d*_*max*_. Using samples from a target new user (*𝒮*_*target*_) as the validation dataset, the error rate of a decision tree was estimated and denoted as *Error*(*Tree, 𝒮*_*target*_). When performing the pruning operation on a node, all its children nodes (and children of children, if any) were removed from the tree. All samples falling into the removed children nodes now fall into their parent nodes, which are the new leaf nodes. The prediction label of the new leaf node after decision tree pruning was defined as the mode of ground truth labels of all validation samples falling into this node. In our bottom-up strategy, we first found the parent nodes of the deepest leaf nodes (denoted as *parentNodes* = *{u*_1_, …, *u*_*i*_, …, *u*_*n*_*}*) with their depth equal to *d*_*max*_. Only if the estimated error rate decreased after pruning these deepest nodes, the pruning operation was performed. The same inspection procedure was repeated on nodes at lower depths until the root node was inspected. The pseudo-code of our pruning strategy is presented in Algorithm 1.

As for grafting, given a set of leaf nodes, *leaf Nodes* = *{u*_1_, …, *u*_*i*_, …, *u*_*n*_*}*, we first constructed a set of validation samples from the new target subject falling into each leaf node *u*_*i*_, denoted as 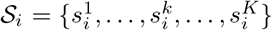. The set of involved ground truth labels of the sample set *𝒮*_*i*_ was defined as 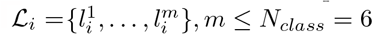. The predicted label of a leaf node *u*_*i*_ was defined as 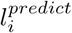 If *m >* 1 (that is, validation samples with more than one label fell into a leaf node *u*_*i*_) or *m* = 1 but 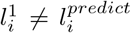, then the decision rule related to the leaf node *u*_*i*_ is not reliable and therefore should be fine-tuned. Concretely, the decision rule related to a leaf node was fine-tuned either by grafting a new decision tree to the leaf node to further purify the sample labels (in the case of *m >* 1), or by directly flipping the predicted label (in the case of *m* = 1 but 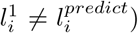. The pseudo-code for our grafting strategy is presented in Algorithm 2.

The decision tree pruning and grafting are opposite operations. Performing either operation is the first step of our model calibration method, and their performances will be compared in our analyses.

#### 2.5.3. Appending Decision Trees

After performing either pruning or grafting on the pre-trained model, we appended an additional 200 decision trees trained merely on data from the new target user. This constructed an RF model with 400 trees. By appending new decision trees, the decision rules of the calibrated RF model consider both the generalised data distribution from a large number of pre-training users and the specific data distribution from the new user.

### 2.6. Baseline Machine Learning Models

Two other machine learning models were implemented to provide baseline performances. First, a user-specific LDA model was trained using data from the new target user. LDA was selected in our work because it is considered as a gold standard for the state-of-the-art EMG pattern recognition models [47, 48, 49]. Second, a user-specific standard RF model was also trained using data from the new target user. The baseline user-specific RF model also consisted of 400 decision trees, the same as the proposed model.

### 2.7. Validation Methods

For the data collected from 20 subjects in experiment 1, we performed an offline leave-one-subject-out cross-validation. For each subject, data from the other 19 subjects were allocated into the pre-training dataset to pre-train an RF model. For the 6 gestures × 10 trials/gesture = 60 trials from the testing subject, only data in the first trial of each gesture were used to calibrate the pre-trained model. To simulate the most challenging validation using minimal calibration data from the testing subject, only data within a 1s duration (randomly segmented out of the total 4s valid duration) of a trial were allocated into the calibration dataset to fine-tune the pre-trained RF. The choice of using only 1s duration of a trial in the calibration dataset is also the same as the counterpart in our subsequent online experiment - experiment 2, where only data within a 1s holding period were available in each calibration trial. For the other two baseline user-specific RF and LDA models, the same calibration dataset was used to train a brand-new user-specific model, and all other configurations were the same. Overall, in our offline validations, we compared the performances of four models, namely, RF calibrated via grafting and appending decision trees, RF calibrated via pruning and appending decision trees, standard RF, and standard LDA.

For our real-time study in experiment 2, an RF model was first pre-trained using all offline data from all 20 subjects of experiment 1. When a new user started the experiment on the first day, data in the 1s holding duration of one trial for each gesture was collected in the calibration session, and used to calibrate the pre-trained RF model. Standard user-specific RF and LDA models were also trained using the same calibration dataset (1 trial per gesture from the new user). In the testing part of experiment 2, the three models, namely RF calibrated via pruning and appending decision trees, the user-specific RF and the user-specific LDA, were implemented and gave their three predictions in each sliding window. Accuracy was evaluated on all predictions in 10 windows within the 1s holding period of a testing trial. On the second day of experiment 2, no model calibration was performed, and the testing section directly started using the same three models trained/calibrated on the first day. A summary of data allocation strategies in both experiments for all models is presented in Table 1.

**Table 1:** Data allocation for different methods. In offline validations, leave-one-subject-out cross-validation was applied, with data from all 20 subjects in experiment 1 used to test all models one by one. In real time experiment, 18 new subjects were used to test all models one by one.

|  |  | Pre-Training |  |  | User-Specific Calibration |  |  |
| --- | --- | --- | --- | --- | --- | --- | --- |
|  |  | Number of Subjects | Number of Repetitions per Gesture per Subject | Signal Duration per Repetition (s) | Number of Subjects | Number of Repetitions per Gesture per Subject | Signal Duration per Repetition (s) |
| Offline Validation | LDA | - | - | - | 1 | 1 | 1 |
|  | RF | - | - | - | 1 | 1 | 1 |
|  | Graft + Append | 19 | 10 | 4 | 1 | 1 | 1 |
|  | Prune + Append | 19 | 10 | 4 | 1 | 1 | 1 |
| Real-Time Experiment | LDA | - | - | - | 1 | 1 | 1 |
|  | RF | - | - | - | 1 | 1 | 1 |
|  | Prune + Append | 20 | 10 | 4 | 1 | 1 | 1 |

Experiment 2 was conducted in an open-loop mode, that is the participants did not see the outcome of the decoders. As such any observed differences between the three decoders is wholly because of the decoding capacity of the model. The experiment was run on a laptop (CPU: 11th Gen Intel(R) Core(TM) i5-1145G7 @ 2.60GHz).

### 2.8. Ablation Experiment

Considering many key components were included in our method, we performed an ablation experiment with each component removed from the processing pipeline, to show the necessity and contribution of each individual component. These key components are: (1) energy descriptors, (2) distribution descriptors, (3) spectrum descriptors, (4) subject-wise feature normalization, (5) pruned decision trees, and (6) appended decision trees. Note that when removing pruned decision trees or appended decision trees, we always keep the same number of total decision trees, namely 400 trees, by doubling the other types of decision trees.

### 2.9. Statistical Analyses

To compare the performances of different models (*≥* 3 groups), the Friedman test was first performed. The Nemenyi post-hoc test, a multi-comparison test to identify pair-wise group differences, was then applied. Significant differences were claimed if *p <* 0.05 was obtained.

## 3. Results

### 3.1. Experiment 1

Results of the first experiment are presented in Figure 3. RF with pruning and appending strategies and RF with grafting and appending strategies yielded an accuracy of 88.3% and 88.2%, respectively, both significantly higher than the outcome of LDA. By definition, pruning leads to smaller models with fewer parameters. Therefore, we selected decision tree pruning rather than grafting in our real-time implementation.

**Figure 3.**
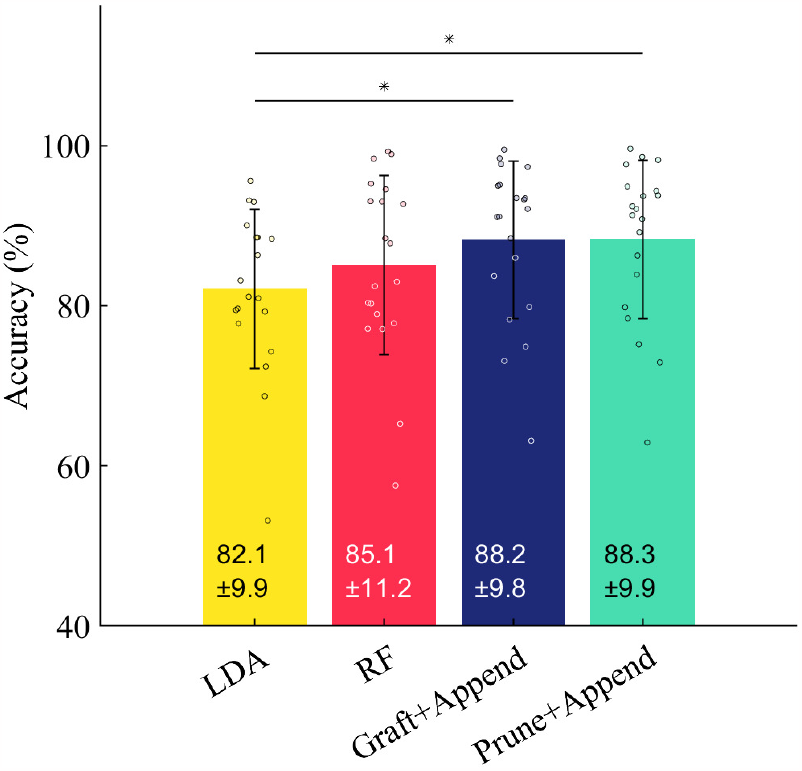
Results in offline validations. Symbol * denotes a significant difference between the two groups.

Additionally, considering the final fine-tuned RF model involves many components, we ran an offline ablation experiment to examine the effect of removing each individual component. Results were presented in Figure 4. Removing each individual component from the overall framework would lead to lower accuracy, indicating the necessity of all these components in the whole framework. Significance was observed when removing the energy features, pruned decision trees, or appended decision trees, demonstrating the important roles of these components.

**Figure 4.**
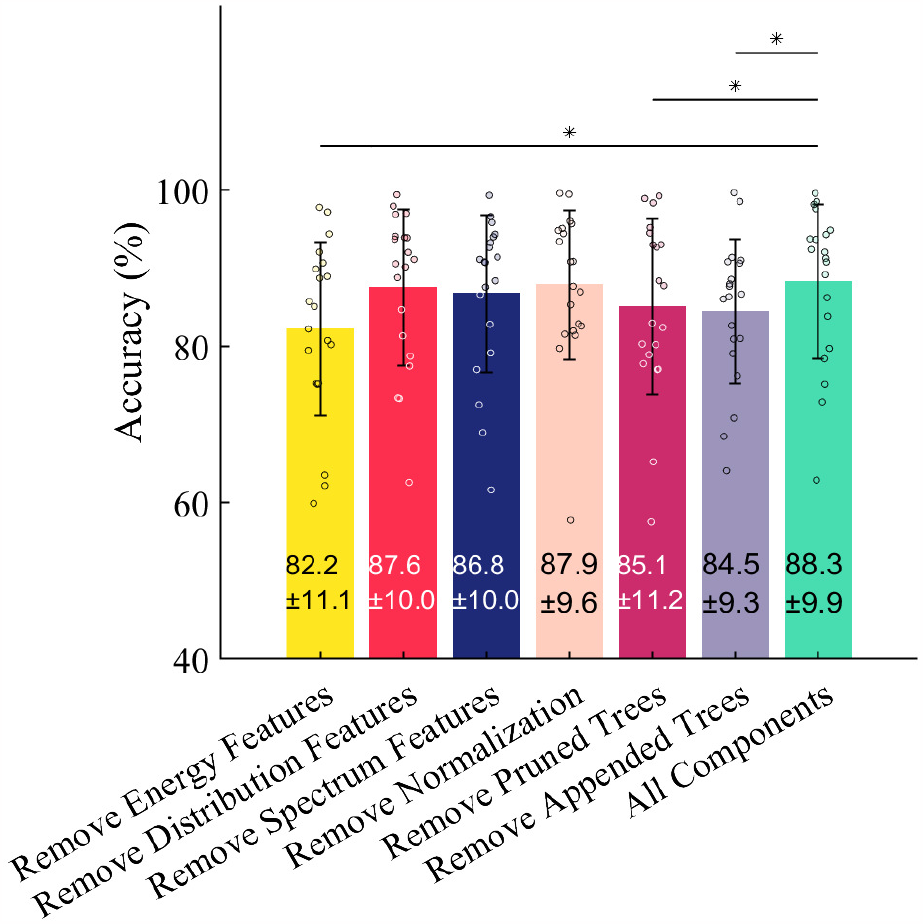
Results in offline ablation experiments. Symbol * denotes a significant difference between the two groups.

### 3.2. Experiment 2

Results of our real-time implementations are presented in Figure 5. On the first day, RF calibrated with pruning and appending decision trees achieved the highest average accuracy of 81.5% but the accuracy improvement over the standard LDA (75.5%) and standard RF (77.6%) is not significant. On the second day, RF calibrated via pruning and appending decision trees significantly outperformed both the standard LDA and the standard RF. This results verifies the the superiority of our method in an inter-day application. The final pre-trained and then fine-tuned RF model involves two groups of decision trees, i.e., the pruned decision trees and the appended decision trees. For the pruned decision trees, because they were pre-trained on a large dataset from a large number of subjects, each decision tree consists of ∼ 925 nodes. For the appended decision trees trained on a small dataset from a new subject, each decision tree comprised ∼ 15 nodes. Accordingly, the whole model consists of 925 × 200 + 15 × 200 = 188, 000 nodes. Four parameters were saved in each node, i.e., the index of the left child (2 bytes), the index of the right child (2 bytes), the index of the selected feature in a node (1 byte) and the threshold of the selected feature (2 bytes), with a size of 7 bytes each node. Therefore, the size of the whole model is 188,000 nodes × 7 bytes/node = 1.26 MB. Other memory usage in the whole processing pipeline is 190.1 *±* 19.2 KB (peak memory).

**Figure 5.**
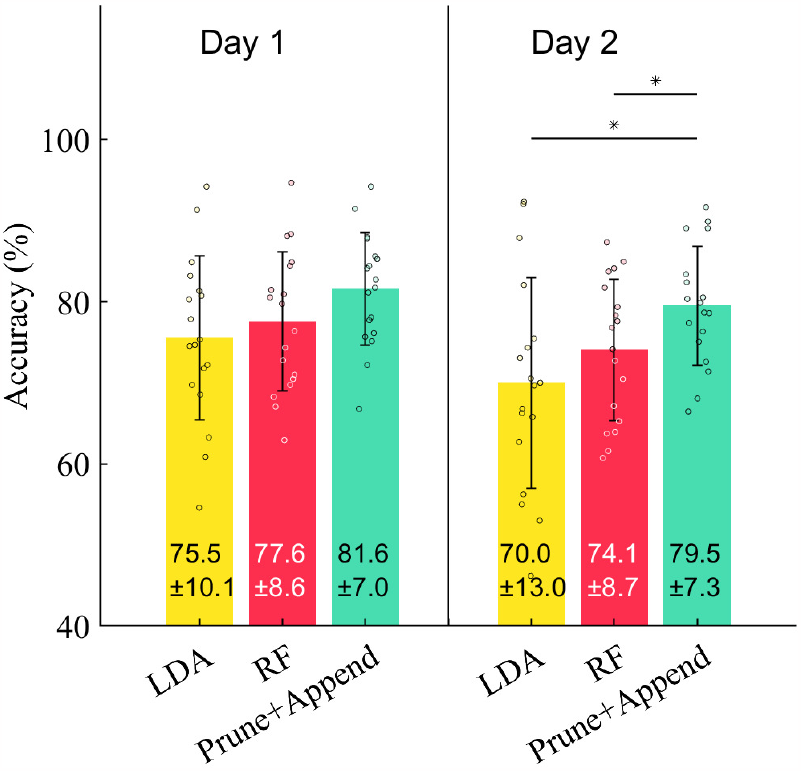
Results in real-time implementation in experiment 2. Symbol * denotes a statistical significance.

## 4. Discussions

We proposed a source-free calibration method for an RF-based myoelectric control model based on pruning and appending decision trees, which outperformed standard LDA and RF models especially in an inter-day study, as presented in Figure 5. The improved performance of RF by pruning and appending decision trees can be explained from two aspects:

- aspect 1: improving the average accuracy of individual decision trees and
- aspect 2: improving the ambiguity (or, diversity) among all decision trees,

which are two important factors determining the overall performance of an ensemble model [50, 51]. In other words, to achieve an excellent ensemble performance, the base learners should be (1) accurate, and (2) different. Accordingly, we evaluated the average accuracy of each individual decision tree and the distribution of their predictions, presented in Figure 6 and Figure 7, respectively. First, as shown in Figure 6, by pruning each pre-trained decision tree, the average accuracy of the pruned decision trees was significantly higher than the pre-trained decision trees, demonstrating that the decision tree pruning operation mainly improves the performance of RF from the above listed aspect 1. Second, as shown in Figure 7, the predictions of the pre-trained trees and pruned trees were similar so their predictions were distributed near each other. This is due to the fact that, the pruning operation was performed on top of each pre-trained tree, so the predictions of pruned trees are more accurate but also quite similar as pre-trained trees. However, the predictions of the appended decision trees were far from the other trees in Figure 7, demonstrating that the appended decision trees could provide different complementary information with the other types of decision trees. The appended trees improve the ambiguity between all decision trees, improving the overall performance from the above listed aspect 2. A high average accuracy and a high ambiguity of decision trees together improve the overall performance of the RF model.

**Figure 6.**
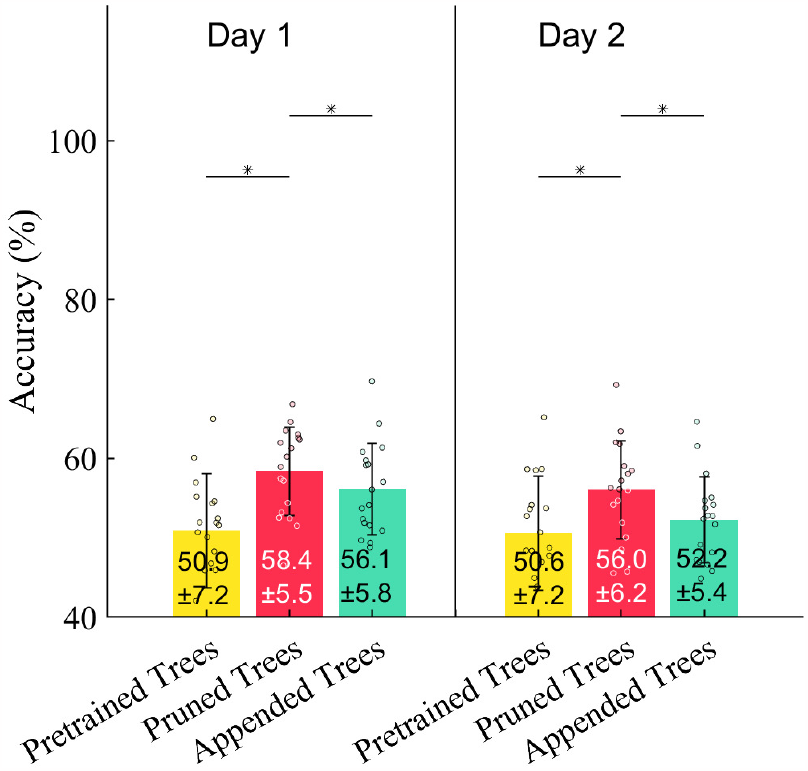
Average accuracy of different decision trees. Results were obtained from each individual decision tree without an ensemble. Symbol * denotes a significant difference between the two groups.

**Figure 7.**
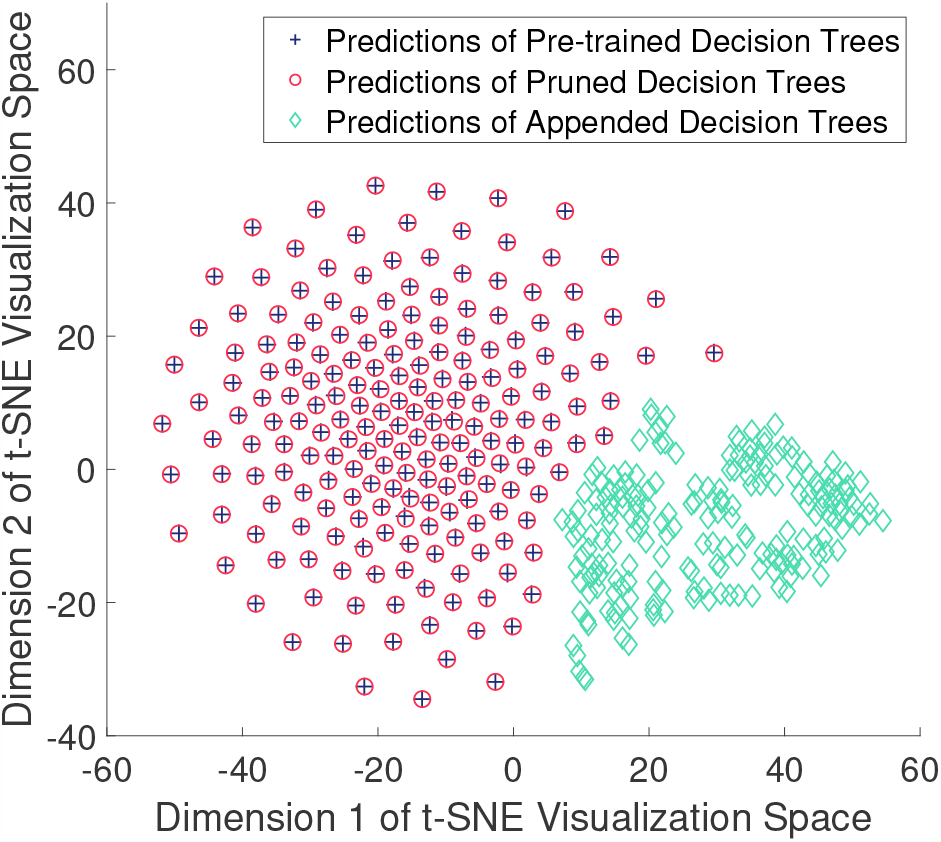
Visualization of predictions of different decision trees in a 2-dimensional space via t-Distributed Stochastic Neighbor Embedding (t-SNE) [52]. Predictions in this figure were drawn from a representative subject on the second day. Predictions on testing samples were used as visualization variables and all decision trees were used as visualization data points. Data from other subjects follow a similar pattern. A single data point represents a decision tree.

We also compared the effectiveness of the grafting and pruning operations on decision trees. Decision tree grafting could further specify the decision rules of a pre-trained decision tree based on the feature distribution of a new user; so that the new decision rules can be better adapted to the new user. By using data from a new user as the validation dataset, pruning could simplify the decision rules. Both grafting and pruning were effective and did not show significantly different performance. We selected decision tree pruning in our real-time implementation because it led to smaller models.

All hyper-parameters were tuned and determined in our offline validations on data collected in experiment 1 and directly used in our real-time implementation in experiment 2 without additional hyper-parameter tuning. One important hyper-parameter is the number of decision trees in the pre-trained RF (200 trees) and appended RF (200 trees). We selected these values because further increasing the number of trees did not contribute to a substantial performance improvement but reduced the computational efficiency. Another hyper-parameter is the number of samples to draw from the pre-training dataset to pre-train each decision tree. Considering the pre-training dataset from multiple subjects is relatively large, drawing the same number of bootstrap samples as the total size of the pre-training dataset would lead to a large decision tree with over-complex decision rules. Therefore, in our work, only 7% samples were randomly drawn from the pre-training dataset via bootstrap to pre-train each decision tree. For user-specific decision trees, considering the size of the calibration dataset from the new target user is small, the number of bootstrap samples to train each user-specific decision tree was the same as the size of the calibration dataset. The tuning of model hyper-parameters in our real-time implementation is pre-determined and completely independent of the data collected in our online experiment.

The significant performance improvement of our method on the second day is another important finding. In inter-day applications of myoelectric control models, the model performance would normally degrade due to the joint effects of multiple factors, e.g. the electrode shift, skin-electrode impedance, etc. By training a new model merely on data from the new user, the model is prone to overfitting on the small dataset from the new user on the first day and degraded performance on the second day. By first pre-training a model and then fine-tuning the model on the new user, the model could adapt to the data of the new user and at the same time retain the generalisation capability learned from a large pre-training dataset. Additionally, the above finding may make even broader contributions, because the pre-training mechanism is expected to improve the inter-day generalisation capability for most machine learning models.

We evaluated the memory requirements and computational efficiency of our method. For memory requirements, we evaluated the size of model parameters and other memory usage in computing (the peak memory). For computational efficiency, we evaluated the computation time of feature extraction and decision-making, respectively. The whole processing pipeline can be divided into two steps, namely, the feature extraction step and the classification step. The computation time in each processing step is presented in Table 2. The feature extraction and classification steps require 2.82 ms and 6.27 ms processing time, respectively, with a total of 9.08 ms for the whole processing pipeline. The processing time of 9.08 ms is short enough for online processing of an EMG segment with a 200 ms window length and 100 ms sliding step. Furthermore, the average model size is 1.26 MB with an additional average memory of 190.1 KB for computations in the whole processing pipeline. These metrics demonstrate the potential of our method for mobile computing scenarios with low-cost computational resources. The computation speed is evaluated using a single CPU without any parallel computing. RF is a model highly suitable for parallel computing because the processing procedures of different decision trees are independent of each other. This property increases the flexibility of the RF model in real-world applications with different computational platforms. The computation speed can be further largely reduced in practice by parallelising different decision trees.

**Table 2:** Computation time in each processing step. No parallel computing was employed.

| Processing Step | Computation Time |
| --- | --- |
| Feature Extraction | 2.8±0.2 ms |
| Classification | 6.2±0.6 ms |
| Total | 9.0±0.7 ms |

Previous studies on transfer learning in myoelectric control mainly focused on deep neural networks [53, 54, 55]. Hoshino et al. [53] achieved a high accuracy of ∼ 95% on 8 hand-wrist gestures, in offline analyses by pre-training and fine-tuning a deep neural network. Côté-Allard et al. [54] achieved a high accuracy of 94.69% on 7 hand-wrist gestures using a similar pre-training and fine-tuning mechanism on deep neural networks. The signal duration for each gesture used in training/validation during the fine-tuning stage in [53] and [54] is 3× and 5×, respectively, of that used in our work. Additionally, the gestures selected in [53, 54] mainly include gross wrist-hand gestures (e.g., wrist extension/flexion) without dexterous control of individual fingers. A very recent work [55] developed a deep convolutional neural network and achieved a high accuracy of 94.94% on 8 hand gestures, using 5 trials (15s signal duration) per gesture for model calibration. The accuracy was substantially dropped to 75.34% using only 1 trial (3s signal duration) per gesture, demonstrating the challenge in achieving a high accuracy with a reduced duration of calibration signals. Additionally, the above accuracy was evaluated on predictions of the whole 3s trials in [55] via majority voting of different windows, while the accuracy reported in our work was evaluated on predictions of 200ms sliding windows facilitating real time hand gesture recognition. In our offline analyses, we achieved an accuracy of 88% on 6 hand gestures using a simple RF model with only 1s calibration data per gesture. Overall, the proposed calibration method provide a brand new transfer learning solution for EMG-based hand gesture recognition using simple, explainable and easily parallelisable RF models.

In our work, we evaluated the effectiveness of RF model calibration on EMG data collected from subjects with intact upper-limbs, proving the high potentials of our method in general human-machine interaction applications. As for applications such as prosthetic control, despite of the variability between EMG of limb-intact subjects and amputees, certain common EMG patterns still remain. A previous study has demonstrated that machine learning models can still learn useful knowledge from sEMG signals of intact subjects, and then help improve model performances on amputees via transfer learning [56]. A very recent study [57] likewise achieved improved performance (improved from 73.4% to 78.1%) on amputees by leveraging knowledge learned from intact subjects. Therefore, the proposed RF-based transfer learning method may also contribute to more applications related to myoelectric control in follow-up studies.

Our work is an initial attempt to demonstrate the superiority of pre-trained and fine-tuned RF over the standard RF in myoelectric control applications. RF has not been considered as one of the most powerful models in myoelectric control. Therefore, little attention has been paid to RF models and there are very few attempts on advanced algorithms for improved RF models [30]. Here we demonstrated the high potential of RF models empowered by several key components such as decision tree pre-training, pruning and appending. Our very recent study [29] also demonstrated the high explainability and robustness of decision tree-based models in myoelectric control. We hope these attempts can attract renewed attention on the efficient, explainable, and powerful decision tree-based models within the myoelectric control community, and expect more interesting and inspiring properties of RF to be discovered in follow-up studies.

## 5. Conclusions

We proposed a myoelectric control model based on random forest, using only 1s of EMG data per gesture from the new user to fine-tune a pre-trained model. Both offline and real-time experiments showed the superiority of our method compared to the standard RF and LDA models. In our inter-day study, our model trained with the data from the first day yielded a significantly higher accuracy compared to other models, without any calibration needed on the second day. Further evaluations also demonstrate the low memory requirements and high computational efficiency of our method.

## References

[1] Nan Zhao, Bolun Zhao, Gencai Shen, Chunpeng Jiang, Zhuangzhuang Wang, Zude Lin, Lanshu Zhou, and Jingquan Liu. A robust hd-semg sensor suitable for convenient acquisition of muscle activity in clinical post-stroke dysphagia. Journal of Neural Engineering, 20(1):016018, jan 2023.

[2] Lahiru N Wimalasena, Jonas F Braun, Mohammad Reza Keshtkaran, David Hofmann, Juan Álvaro Gallego, Cristiano Alessandro, Matthew C Tresch, Lee E Miller, and Chethan Pandarinath. Estimating muscle activation from emg using deep learning-based dynamical systems models. Journal of Neural Engineering, 19(3):036013, may 2022.

[3] Yue Wen, Sangjoon J Kim, Simon Avrillon, Jackson T Levine, François Hug, and José L Pons. Toward a generalizable deep cnn for neural drive estimation across muscles and participants. Journal of Neural Engineering, 20(1):016006, jan 2023.

[4] Xuan Zhang, L. Wu, Xu Zhang, Xiang Chen, Chang Li, and Xun Chen. Multi-source domain generalization and adaptation toward cross-subject myoelectric pattern recognition. Journal of Neural Engineering, 20(1):016050, feb 2023.

[5] Daniela Souza de Oliveira, Andrea Casolo, Thomas G Balshaw, Sumiaki Maeo, Marcel Bahia Lanza, Neil R W Martin, Nicola Maffulli, Thomas Mehari Kinfe, Bjoern M Eskofier, Jonathan P Folland, Dario Farina, and Alessandro Del Vecchio. Neural decoding from surface high-density emg signals: influence of anatomy and synchronization on the number of identified motor units. Journal of Neural Engineering, 19(4):046029, aug 2022.

[6] Yanan Diao, Qiangqiang Chen, Yan Liu, Linjie He, Yue Sun, Xiangxin Li, Yumin Chen, Guanglin Li, and Guoru Zhao. A fuzzy granular logistic regression algorithm for semg-based cross-individual prosthetic hand gesture classification. Journal of Neural Engineering, 20(2):026029, apr 2023.

[7] Angkoon Phinyomark, Pornchai Phukpattaranont, and Chusak Limsakul. Feature reduction and selection for emg signal classification. Expert Systems with Applications, 39(8):7420–7431, 2012.

[8] Ali H. Al-Timemy, Guido Bugmann, Javier Escudero, and Nicholas Outram. Classification of finger movements for the dexterous hand prosthesis control with surface electromyography. IEEE Journal of Biomedical and Health Informatics, 17(3):608–618, 2013.

[9] Farshad Khadivar, Vincent Mendez, Carolina Correia, Iason Batzianoulis, Aude Billard, and Silvestro Micera. Emg-driven shared human-robot compliant control for in-hand object manipulation in hand prostheses. Journal of Neural Engineering, 19(6):066024, ec 2022.

[10] Pranav Mamidanna, Jakob L Dideriksen, and Strahinja Dosen. Estimating speed-accuracy trade-offs to evaluate and understand closed-loop prosthesis interfaces. Journal of Neural Engineering, 19(5):056012, sep 2022.

[11] Federica Barberi, Francesco Iberite, Eugenio Anselmino, Pericle Randi, Rinaldo Sacchetti, Emanuele Gruppioni, Alberto Mazzoni, and Silvestro Micera. Early decoding of walking tasks with minimal set of emg channels. Journal of Neural Engineering, 20(2):026038, apr 2023.

[12] Philip P Vu, Alex K Vaskov, Christina Lee, Ritvik R Jillala, Dylan M Wallace, Alicia J Davis, Theodore A Kung, Stephen W P Kemp, Deanna H Gates, Cynthia A Chestek, and Paul S Cederna. Long-term upper-extremity prosthetic control using regenerative peripheral nerve interfaces and implanted emg electrodes. Journal of Neural Engineering, 20(2):026039, apr 2023.

[13] Md Abdur Rahim and Jungpil Shin. Hand movement activity-based character input system on a virtual keyboard. Electronics, 9(5), 2020.

[14] Md. Rokib Raihan and Mohiuddin Ahmad. Developing wearable human–computer interfacing system based on emg and gyro for amputees. In Mohiuddin Ahmad, Mohammad Shorif Uddin, and Yeong Min Jang, editors, Proceedings of International Conference on Information and Communication Technology for Development, pages 291–304, Singapore, 2023. Springer Nature Singapore.

[15] Feiyun Xiao, Jingsong Mu, Jieping Lu, Guangxu Dong, and Yong Wang. Real-time modeling and feature extraction method of surface electromyography signal for hand movement classification based on oscillatory theory. Journal of Neural Engineering, 19(2):026011, mar 2022.

[16] Asim Waris, Imran K. Niazi, Mohsin Jamil, Kevin Englehart, Winnie Jensen, and Ernest Nlandu Kamavuako. Multiday evaluation of techniques for emg-based classification of hand motions. IEEE Journal of Biomedical and Health Informatics, 23(4):1526–1534, 2019.

[17] Katarzyna Szymaniak, Agamemnon Krasoulis, and Kianoush Nazarpour. Recalibration of myoelectric control with active learning. Frontiers in Neurorobotics, 16, 2022.

[18] Abolfazl Farahani, Sahar Voghoei, Khaled Rasheed, and Hamid R. Arabnia. A brief review of domain adaptation. In Robert Stahlbock, Gary M. Weiss, Mahmoud Abou-Nasr, Cheng-Ying Yang, Hamid R. Arabnia, and Leonidas Deligiannidis, editors, Advances in Data Science and Information Engineering, pages 877–894, Cham, 2021. Springer International Publishing.

[19] Marina M.-C. Vidovic, Han-Jeong Hwang, Sebastian Amsüss, Janne M. Hahne, Dario Farina, and Klaus-Robert Müller. Improving the robustness of myoelectric pattern recognition for upper limb prostheses by covariate shift adaptation. IEEE Transactions on Neural Systems and Rehabilitation Engineering, 24(9):961–970, 2016.

[20] Jinqiang Wang, Dianguo Cao, Yang Li, Jiashuai Wang, and Yuqiang Wu. Multi-user motion recognition using semg via discriminative canonical correlation analysis and adaptive dimensionality reduction. Frontiers in Neurorobotics, 16, 2022.

[21] Bo Xue, L. Wu, Kun Wang, Xu Zhang, Juan Cheng, Xiang Chen, and Xun Chen. Multiuser gesture recognition using semg signals via canonical correlation analysis and optimal transport. Computers in Biology and Medicine, 130:104188, 2021.

[22] Xinyu Jiang, Berj Bardizbanian, Chenyun Dai, Wei Chen, and Edward A. Clancy. Data management for transfer learning approaches to elbow emg-torque modeling. IEEE Transactions on Biomedical Engineering, 68(8):2592–2601, 2021.

[23] Machiel J Zwarts, Gea Drost, and Dick F Stegeman. Recent progress in the diagnostic use of surface emg for neurological diseases. Journal of Electromyography and Kinesiology, 10(5):287–291, 2000.

[24] Ashirbad Pradhan, Jiayuan He, and Ning Jiang. Performance optimization of surface electromyography based biometric sensing system for both verification and identification. IEEE Sensors Journal, 21(19):21718–21729, 2021.

[25] Baolin Peng, Chunyuan Li, Pengcheng He, Michel Galley, and Jianfeng Gao. Instruction tuning with gpt-4, 2023.

[26] Alexander Kirillov, Eric Mintun, Nikhila Ravi, Hanzi Mao, Chloe Rolland, Laura Gustafson, Tete Xiao, Spencer Whitehead, Alexander C. Berg, Wan-Yen Lo, Piotr Dollár, and Ross Girshick. Segment anything, 2023.

[27] Paras Gulati, Qin Hu, and S. Farokh Atashzar. Toward deep generalization of peripheral emg-based human-robot interfacing: A hybrid explainable solution for neurorobotic systems. IEEE Robotics and Automation Letters, 6(2):2650–2657, 2021.

[28] Hyunin Lee, Dongwook Kim, and Yong-Lae Park. Explainable deep learning model for emg-based finger angle estimation using attention. IEEE Transactions on Neural Systems and Rehabilitation Engineering, 30:1877–1886, 2022.

[29] Xinyu Jiang, Kianoush Nazarpour, and Chenyun Dai. Explainable and robust deep forests for emg-force modeling. IEEE Journal of Biomedical and Health Informatics, pages 1–12, 2023.

[30] Tao Zhou, Olatunji Mumini Omisore, Wenjing Du, Lei Wang, and Yuan Zhang. Adapting random forest classifier based on single and multiple features for surface electromyography signal recognition. In 2019 12th International Congress on Image and Signal Processing, BioMedical Engineering and Informatics (CISP-BMEI), pages 1–6, 2019.

[31] Yanjun Qi. Random Forest for Bioinformatics, pages 307–323. Springer US, Boston, MA, 2012.

[32] Trevor Hastie, Jerome Friedman, and Robert Tibshirani. The Elements of Statistical Learning: Data Mining, Inference, and Prediction. Springer, Berlin, 2009.

[33] Gérard Biau and Erwan Scornet. A random forest guided tour. TEST, 25(2):197–227, Jun 2016.

[34] Riccardo Guidotti, Anna Monreale, Salvatore Ruggieri, Franco Turini, Fosca Giannotti, and Dino Pedreschi. A survey of methods for explaining black box models. ACM Comput. Surv., 51(5), aug 2018.

[35] John Mingers. An empirical comparison of pruning methods for decision tree induction. Machine learning, 4:227–243, 1989.

[36] Noam Segev, Maayan Harel, Shie Mannor, Koby Crammer, and Ran El-Yaniv. Learn on source, refine on target: A model transfer learning framework with random forests. IEEE Transactions on Pattern Analysis and Machine Intelligence, 39(9):1811–1824, 2017.

[37] Geoffrey I. Webb. Decision tree grafting from the all-tests-but-one partition. In 16th International Joint Conference on Artificial Intelligence Proceedings, volume 2 of IJCAI International Joint Conference on Artificial Intelligence, pages 702–707, December 1999. International Joint Conference on Artificial Intelligence 1999, IJCAI 1999 ; Conference date: 31-07-1999 Through 06-08-1999.

[38] B. Hudgins, P. Parker, and R.N. Scott. A new strategy for multifunction myoelectric control. IEEE Transactions on Biomedical Engineering, 40(1):82–94, 1993.

[39] Fuqiang Ye, Bibo Yang, Chingyi Nam, Yunong Xie, Fei Chen, and Xiaoling Hu. A data-driven investigation on surface electromyography based clinical assessment in chronic stroke. Frontiers in Neurorobotics, 15, 2021.

[40] K. Englehart and B. Hudgins. A robust, real-time control scheme for multifunction myoelectric control. IEEE Transactions on Biomedical Engineering, 50(7):848–854, 2003.

[41] Egon L. van den Broek, Marleen H. Schut, Joyce H. D. M. Westerink, Jan van Herk, and Kees Tuinenbreijer. Computing emotion awareness through facial electromyography. In Thomas S. Huang, Nicu Sebe, Michael S. Lew, Vladimir Pavlović, Mathias Kölsch, Aphrodite Galata, and Branislav Kisačanin, editors, Computer Vision in Human-Computer Interaction, pages 52–63, Berlin, Heidelberg, 2006. Springer Berlin Heidelberg.

[42] Kianoush Nazarpour, Ahmad R. Sharafat, and S. Mohammad P. Firoozabadi. Application of higher order statistics to surface electromyogram signal classification. IEEE Transactions on Biomedical Engineering, 54(10):1762–1769, 2007.

[43] Rami N. Khushaba, Ahmed Al-Ani, and Adel Al-Jumaily. Orthogonal fuzzy neighborhood discriminant analysis for multifunction myoelectric hand control. IEEE Transactions on Biomedical Engineering, 57(6):1410–1419, 2010.

[44] Angkoon Phinyomark, Pornchai Phukpattaranont, and Chusak Limsakul. Feature reduction and selection for emg signal classification. Expert Systems with Applications, 39(8):7420–7431, 2012.

[45] F. Esposito, D. Malerba, G. Semeraro, and J. Kay. A comparative analysis of methods for pruning decision trees. IEEE Transactions on Pattern Analysis and Machine Intelligence, 19(5):476–491, 1997.

[46] Floriana Esposito, Donato Malerba, and Giovanni Semeraro. Simplifying decision trees by pruning and grafting: New results (extended abstract). In Nada Lavrac and Stefan Wrobel, editors, Machine Learning: ECML-95, pages 287–290, Berlin, Heidelberg, 1995. Springer Berlin Heidelberg.

[47] Martyna Stachaczyk. Decoding peripheral neural correlates of dexterous movements. 2022.

[48] Ning Jiang, Thomas Lorrain, and Dario Farina. A state-based, proportional myoelectric control method: online validation and comparison with the clinical state-of-the-art. Journal of NeuroEngineering and Rehabilitation, 11(1):110, Jul 2014.

[49] Alexander E. Olsson, Nebojša Malešević, Anders Björkman, and Christian Antfolk. Learning regularized representations of categorically labelled surface emg enables simultaneous and proportional myoelectric control. Journal of NeuroEngineering and Rehabilitation, 18(1):35, Feb 2021.

[50] Anders Krogh and Jesper Vedelsby. Neural network ensembles, cross validation, and active learning. In G. Tesauro, D. Touretzky, and T. Leen, editors, Advances in Neural Information Processing Systems, volume 7. MIT Press, 1994.

[51] Zhi-Hua Zhou and Ji Feng. Deep forest. National Science Review, 6(1):74–86, 10 2018.

[52] Laurens van der Maaten and Geoffrey Hinton. Visualizing Data using t-SNE. Journal of Machine Learning Research, 9(Nov):2579–2605, 2008.

[53] Takayuki Hoshino, Suguru Kanoga, Masashi Tsubaki, and Atsushi Aoyama. Comparing subject-to-subject transfer learning methods in surface electromyogram-based motion recognition with shallow and deep classifiers. Neurocomputing, 489:599–612, 2022.

[54] Ulysse Côté-Allard, Cheikh Latyr Fall, Alexandre Drouin, Alexandre Campeau-Lecours, Clément Gosselin, Kyrre Glette, François Laviolette, and Benoit Gosselin. Deep learning for electromyographic hand gesture signal classification using transfer learning. IEEE Transactions on Neural Systems and Rehabilitation Engineering, 27(4):760–771, 2019.

[55] Md. Rabiul Islam, Daniel Massicotte, Philippe Y. Massicotte, and Wei-Ping Zhu. Surface emg-based inter-session/inter-subject gesture recognition by leveraging lightweight all-convnet and transfer learning, 2023.

[56] Jinghua Fan, Mingzhe Jiang, Chuang Lin, Gloria Li, Jinan Fiaidhi, Chenfei Ma, and Wanqing Wu. Improving semg-based motion intention recognition for upper-limb amputees using transfer learning. Neural Computing and Applications, 35(22):16101–16111, Aug 2023.

[57] Chuang Lin, Xinyue Niu, Jun Zhang, and Xianping Fu. Improving motion intention recognition for trans-radial amputees based on semg and transfer learning. Applied Sciences, 13(19), 2023.

